# Phosphoribulokinase abundance is not limiting the Calvin-Benson-Bassham cycle in *Chlamydomonas reinhardtii*

**DOI:** 10.1101/2023.05.10.540127

**Authors:** Nicolas D. Boisset, Giusi Favoino, Maria Meloni, Lucile Jomat, Corinne Cassier-Chauvat, Mirko Zaffagnini, Stéphane D. Lemaire, Pierre Crozet

## Abstract

Improving photosynthetic efficiency in plants and microalgae is of utmost importance to support the growing world population and to enable the bioproduction of energy and chemicals. Limitations in photosynthetic light conversion efficiency can be directly attributed to kinetic bottlenecks within the Calvin-Benson-Bassham cycle (CBBC) responsible for carbon fixation. A better understanding of these bottlenecks *in vivo* is crucial to overcome these limiting factors through bio-engineering. The present study is focused on the analysis of phosphoribulokinase (PRK) in the unicellular green alga *Chlamydomonas reinhardtii*. We have characterized a PRK knock-out mutant strain and showed that in the absence of PRK, Chlamydomonas cannot grow photoautotrophically while functional complementation with a synthetic construct allowed restoration of photoautotrophy. Nevertheless, using standard genetic elements, the expression of PRK was limited to 40% of the reference level in complemented strains and could not restore normal growth in photoautotrophic conditions suggesting that the CBBC is limited. We were subsequently able to overcome this initial limitation by improving the design of the transcriptional unit expressing PRK using diverse combinations of DNA parts including PRK endogenous promoter and introns. This enabled us to obtain strains with PRK levels comparable to the reference strain and even overexpressing strains. A collection of strains with PRK levels between 16% and 250% of WT PRK levels was generated and characterized. Immunoblot and growth assays revealed that a PRK content of ≈86% is sufficient to fully restore photoautotrophic growth. This result suggests that PRK is present in moderate excess in Chlamydomonas. Consistently, the overexpression of PRK did not increase photosynthetic growth indicating that that the endogenous level of PRK in Chlamydomonas is not limiting the Calvin-Benson-Bassham cycle under optimal conditions.

## 1. Introduction

The irreversible depletion of traditional sources of fossil fuels coupled with the accumulation of greenhouse gases produced by their combustion has created an urgent need to develop alternative forms of eco-responsible processes for large scale CO_2_ sequestration and new sources of reduced carbon for the production of fuels and chemicals needed by our society (Eckardt et al., 2023). Photoautotrophic microorganisms such as green microalgae and cyanobacteria, are regarded as promising platforms for the development of innovative concepts based on their inherent ability to fix CO_2_, thereby producing various organic molecules *via* a sunlight-driven and sustainable process. Simultaneously, the global crop production needs to double by 2050 to meet the demand of a growing population, especially considering the use of arable lands to feed bio-refineries, deleterious effects of climate change, and continuous erosion of agricultural land (Godfray et al., 2010; Tilman et al., 2011; Ort et al., 2015; Simkin et al., 2019). The remarkable gains in productivity of the Green Revolution of the late 20th century have largely been achieved by increasing the light capture efficiency and the harvest index (*i*.*e*. the fraction of biomass that is captured in the harvested part); but these two factors approach their practical limits (Long et al., 2006). Improved solar energy conversion efficiency (*i*.*e*., photosynthetic efficiency) has so far played little role in improving yield potential, yet photosynthesis is the only determinant that is not close to its biological limits (Zhu et al., 2010).

In order to face the challenge of improving photosynthetic efficiency, new methodologies are required to allow success (Erb and Zarzycki, 2016; Kubis and Bar-Even, 2019; da Fonseca-Pereira et al., 2022). Biology is currently facing a revolution through its transition from analytic to synthetic biology approaches. The rise of green synthetic biology offers the potential to tackle the challenge of improving photosynthetic efficiency through engineering of microalgae, cyanobacteria and plants. Different strategies have shown the potential of synthetic approaches aimed at improving carbon fixation through rewiring of photorespiration (Trudeau et al., 2018; South et al., 2019; Wang et al., 2020; Roell et al., 2021; Jin et al., 2023), engineering of carbon concentration mechanisms (Long et al., 2016; Long et al., 2018; Mackinder, 2018; Atkinson et al., 2020; Hennacy and Jonikas, 2020; Adler et al., 2022) or cutting respiratory carbon losses (Long et al., 2015; Amthor et al., 2019; Garcia et al., 2023). One of the strategies with the highest potential consists in engineering redesigned or synthetic CO_2_ fixation pathways (Long et al., 2015). In the light, the photosynthetic electron transfer (PET) chain produces both energy (ATP) and reducing power (NADPH), which are mainly used by the Calvin-Benson-Bassham cycle (CBBC) to fix atmospheric CO_2_ thereby generating triose phosphate as an immediate product (Vecchi et al., 2020; Raines, 2022). The CBBC comprises 11 enzymes catalyzing 13 reactions and three enzymes are specific to the cycle: ribulose-1,5-bisphosphate carboxylase oxygenase (Rubisco), sedoheptulose-1,7-bisphosphatase (SBPase) and phosphoribulokinase (PRK) (Michelet et al., 2013; Le Moigne et al., 2023; Meloni et al., 2023).

Limitations in photosynthetic light conversion efficiency can be directly attributed to kinetic bottlenecks within the CBBC (Stitt et al., 2010; Raines, 2011). At moderate to high light intensities, the slow turnover of the CBBC leads to overreduction of the PET and results in the dissipation of the excess energy as heat, fluorescence or increased production of reactive oxygen species (Marcus et al., 2011; Wobbe and Remacle, 2015). Therefore, improving the CBBC turnover through synthetic biology approaches is a major avenue to enhance photosynthetic efficiency and increase production of biomass and chemicals. An innovative approach would be to replace the natural CBBC by a more efficient artificial synthetic carbon fixation pathway. Several cycles, which are theoretically more efficient than the CBBC have been proposed or even tested *in vitro* but remain to be validated *in vivo* (Bar-Even et al., 2010; Schwander et al., 2016; Bar-Even, 2018; Gleizer et al., 2019; Satanowski et al., 2020).

The green microalga *Chlamydomonas reinhardtii* (hereafter Chlamydomonas) has a photosynthetic apparatus very similar to that of land plants, and our long-term knowledge of its genetics and physiology make it a good model system to study the CBBC (Salome and Merchant, 2019; Le Moigne et al., 2023). Notably, it is able to grow fast either photoautotrophically or heterotrophically in the presence of a reduced carbon source (acetate), allowing growth of photosynthetic mutants. Moreover, Chlamydomonas is a suitable chassis for synthetic biology approaches (Vavitsas et al., 2019). Diverse key enabling technologies are available including a Chlamydomonas modular cloning toolkit comprising 120 bricks allowing fast and easy generation of any multigenic assembly (Crozet et al., 2018; de Carpentier et al., 2020), CRISPR/Cas9 genome editing techniques (Shin et al., 2016; Ferenczi et al., 2017; Greiner et al., 2017; Kim et al., 2020), and a collection of mapped insertional mutants covering 83% of the nuclear genes (Li et al., 2019). Hence, Chlamydomonas appears as a very good chassis to explore *in vivo* the synthetic redesign of the CBBC and its potential to improve biomass and bioproduct production.

Under saturating light, the CBBC is co-limited by the low catalytic efficiency of Rubisco-dependent carboxylation and by the capacity for regeneration of the Rubisco substrate ribulose-1,5-bisphosphate (RuBP). Strategies for rationally redesigning Rubisco in order to improve its catalytic features or its substrate specificity have generally failed (Kubis and Bar-Even, 2019), although some directed evolution or overexpression strategies were shown to have some potential in several species (Iwaki et al., 2006; Durao et al., 2015; Liang and Lindblad, 2016; 2017; Salesse-Smith et al., 2018; Yoon et al., 2020). Other enzymes also limit CBBC turnover as suggested by modeling and metabolic flux control analyses (Stitt and Schulze, 1994; Raines, 2003; Zhu et al., 2007; Janasch et al., 2019; Raines, 2022). The overexpression of SBPase was shown to improve biomass production and carbon fixation in numerous species (Lefebvre et al., 2005; Tamoi et al., 2006; Rosenthal et al., 2011; Fang et al., 2012; Ding et al., 2016; Liang and Lindblad, 2016; Driever et al., 2017). Similarly, in *Chlamydomonas*, a 3-fold increase of SBPase content increased both photosynthetic rate and growth under high-light and high CO_2_ (Hammel et al., 2020). Besides Rubisco and SBPase, the limitations imposed by the third CBBC specific enzyme, PRK, remain largely unexplored. This enzyme catalyzes the phosphorylation of ribulose-5-phosphate into RuBP, the substrate of Rubisco. The biochemical properties of PRK have been extensively studied *in vitro* (Michelet et al., 2013; Le Moigne et al., 2023; Meloni et al., 2023). These studies showed that, *in vitro*, this enzyme is tightly regulated by light through oxidoreduction of specific disulfides by thioredoxins. PRK activity is regulated both autonomously through reduction of an intramolecular disulfide by thioredoxin that couples PRK activity to light intensity, and non-autonomously through formation in the dark of an inhibitory supramolecular complex with the CP12 protein and the CBBC enzyme glyceraldehyde-3-phosphate dehydrogenase (Gurrieri et al., 2021; Gurrieri et al., 2023). The formation and dissociation of the complex is under the control of thioredoxins through reduction of disulfides on CP12 and PRK (Marri et al., 2008; Marri et al., 2009; Thieulin-Pardo et al., 2015). The structure of oxygenic photosynthetic PRK has been solved for Chlamydomonas, *Arabidopsis thaliana*, and *Synechococcus elongatus* (Gurrieri et al., 2019; Wilson et al., 2019; Yu et al., 2020), as well as the structure of the GAPDH-CP12-PRK complex (McFarlane et al., 2019; Yu et al., 2020). Altering the level of PRK in tobacco using antisense RNA approaches revealed that only plants with a PRK activity below 15% showed decreased carbon fixation (Paul et al., 1995; Banks et al., 1999). This suggests that PRK content might not be limiting the CBBC in tobacco although this was not confirmed by measuring PRK protein content or by overexpression of PRK. In Chlamydomonas, a strain lacking PRK activity was reported to be inefficient for photosynthetic carbon fixation (Moll and Levine, 1970) but this mutant was later found to revert spontaneously precluding its use for engineering approaches ((Smith, 2000) and personal observation of the authors).

In the present study, we characterized a PRK knock-out mutant strain and demonstrated that, Chlamydomonas cannot grow photoautotrophically in the absence of PRK, while functional complementation with a synthetic construct allowed the restoration of photoautotrophy. However, using standard genetic elements, PRK expression was limited to 16-40% of the reference level in complemented strains. By improving the design of the transcriptional unit expressing PRK using various combinations of DNA parts, including the endogenous PRK promoter and introns, we overcame this initial limitation. This enabled us to obtain strains with PRK levels comparable to the reference strain and even overexpressing strains. We generated and characterized a collection of strains with PRK levels between 16% and 250% of WT PRK levels.

## 2. Materials and Methods

### 2.1. Strains, media and growth conditions

The strains used in this study originate from the CLiP library (Li et al., 2019): the reference strain (CC-4533) and ΔPRK (LMJ.RY0402.119555), both obtained from the [Chlamydomonas Resource Center](http://www.chlamycollection.org/). Chlamydomonas cells were grown on agar plates or liquid medium, using Tris-acetate-phosphate (TAP) medium (Gorman and Levine, 1965) or High salt medium (HSM) (Sueoka, 1960) at 25°C, under continuous light (40-60 μmol photons m^-2^ s^-1^) or dark (in particular for the ΔPRK strain), and shaking for liquid cultures (130 rpm). Antibiotics used were hygromycin B (10 μg/mL) and paromomycin (15 μg/mL). Growth analyses were performed in the Algem® labscale double photobioreactor systems (Algenuity, Stewartby, UK) for large volume cultures (400 mL) or in the Algem® HT-24 (Algenuity) photobioreactor for small volume cultures (25 mL). All chemicals were obtained from Sigma-Aldrich unless otherwise specified.

### 2.2. Plasmid construction

Protein and nucleic acid designs were performed *in silico* using Serial Cloner 2.6.1 software. All recipient plasmids are derived from the MoClo original toolkit (Weber et al., 2011).

Phosphoribulokinase PRK cDNA sequence (Cre12.g554800; Uniprot accession P19824) was obtained by PCR on reverse translated mRNA extracts from a D66 strain (Schnell and Lefebvre, 1993) using primers TTGAAGACTTAATGGCTTTCACTATGCGCGC and TTGAAGACAACGAACCCACGGGCACAACGTCC. The resulting PRK coding sequence was designed for the position B3-B4 of the Chlamydomonas MoClo toolkit (Crozet et al., 2018), and cloned into the plasmid pAGM1287 (Weber et al., 2011). Two other constructs were obtained by PCR on genomic DNA of CC-4533 strain using the following primers: TTGAAGACTTCTCAGGAGGCCCTGGGCTTTAGCCCC and AAGAAGACAACTCGAGTACATGATGCATGTAACAGCAGCAATGAT for the PRK promoter, TTGAAGACTTCTCATACTGCGTCTTGGGTCGGTGCGCT and CAGAAGACAACTCGCATTGGTTGCTAACAGCTCGACGC for the PRK 5’UTR, and TTGAAGACTTCTCAAATGGCTTTCACTATGCGCGC and TTGAAGACTTCTCAAATGGCTTTCACTATGCGCGC for the PRK CDS with introns.

Each part was cloned into the plasmid pAGM9121 (Patron et al., 2015). The level 1 plasmids were built in pICH47742 with the endogenous promoter and 5’UTR of PRK and the 3’UTR/Terminator of PSAD controlling the expression of the PRK CDS with or without its endogenous introns. Level M plasmids were built in pAGM8031 combining p1-013 (hygromycin resistance gene) (de Carpentier et al., 2020) with the PRK transcriptional unit resulting in pCMM-24 (P_PSAD_-PRK_CDS_), pCMM-25 (P_PRK_- PRK_CDS_) and pCMM-26 (P_PRK_-PRK_(i)_).

### 2.3. Chlamydomonas transformation

Transformations were performed as previously described (Crozet et al., 2018), using 55 fmol of purified cassette after *Bbs*I-HF (for photoautotrophy screening with p1 plasmids) or *Bsa*I-HF (for antibiotic screening with pM plasmids) digestion (New England Biolabs) of the corresponding plasmid. Transformants were selected on HSM-agar medium or TAP-agar containing hygromycin B (20 mg/L), Plates and transformants were analyzed after 5 to 7 days of growth in medium light (50 μmol photons m^-2^ s^-1^).

### 2.4. Chlamydomonas genotyping

Cells were grown in TAP medium up to 4-5 × 10^6^ cells mL^−1^, harvested by centrifugation at 2500 *g* for 5 min at room temperature (RT), and lysed in 400 μL of extraction buffer (0.2 M Tris-HCl pH 7.5, 200 mM NaCl; 25 mM EDTA; 0.5% SDS) for 10 min at 37°C under agitation (1400 rpm). After centrifugation at 17000 *g* for 3 min at RT, the genomic DNA contained in the supernatant was precipitated with one volume of isopropanol for 10 min at room temperature and collected by centrifugation at 17000 *g* for 10 min at RT. The pellet DNA was then washed with 70% ethanol, spinned (17000 *g* for 3 min at RT) and the pellet was air-dried prior to resuspension in water. PCR was performed using the Quick-LoadR©Taq2×Master Mix (New England Biolabs) according to the manufacturer’s instructions. Primers used were Plex5: AAGGACGCTGACATG, Plm1: CCTGATGGATGGTTC, PLPSAD: TTGAAGACAATCATCTCAATGGGTGTG and Plen3U: AGGTGCCAAAGCAAC.

### 2.5. Protein extraction

Cells were grown in TAP medium up to 4-5 × 10^6^ cells mL^−1^, harvested by centrifugation at 5000 *g* for 10 min at 4°C, resuspended in 500 μL of Buffer B (30 mM Tris-HCl pH 7.9, 0.5 mM EDTA, antiprotease complete tablets (Roche)), and lysed twice by using glass beads and vortexing (30 sec vortex, 1 min on ice). The total extract was then clarified by centrifugation (2×10 min at 21000 *g*) and the concentration of the total soluble protein was determined by BCA Protein Assay using bovine serum albumin (BSA) as standard.

### 2.6. Western blot

Total soluble proteins were analyzed by western blotting with a custom rabbit polyclonal primary antibody raised against Chlamydomonas PRK (Covalab, Bron, France) subsequently detected by secondary anti-rabbit antibody coupled to horseradish peroxidase (Sigma-Aldrich reference A9169, Saint Louis, USA). Detection was done with commercial ECL peroxidase assay (GE Healthcare, Chicago IL USA) with a Chemidoc (Bio-Rad, Hercules CA USA).

### 2.7. Spot tests

Cells were grown until exponential phase (2-6 × 10^6^ cells mL^−1^) and serial dilutions in TAP or HSM media were made. The dilution was spotted (10 μL) onto TAP and HSM agar plates at different light intensities. The plates were then scanned after 7 days using a Perfection V800 Photo scanner (Epson). This analysis was automatized using an Opentrons OT-2.

### 2.8. Growth analysis in photobioreactor

Growth analyses were performed using the Algem® labscale double photobioreactor system (Algenuity, Stewartby, United Kingdom) under continuous light (100 μmol photons m^-2^ s^-1^) and 120 rpm agitation in TAP or high salt medium (HSM). The absorbance at 740 nm was recorded every 10 min using the built-in sensor. The maximal growth rate was determined as the maximal slope of the growth curve (ΔAbs/Δtime). Growth curves obtained with the AlgemHT24 photobioreactor were performed in autotrophic conditions and with three different light intensities, *i*.*e*. low light at 38 μmol.m^-2^.s^-1^, medium light at 100 μmol.m^-2^.s^-1^, and high light at 300 μmol.m^-2^.s^-1^. The resulting growth curves were analyzed using GraphPad Prism software, and in particular a fit was made with the Gompertz growth equation (Y = YM*(Y0/YM)^(exp(-K*X))) to obtain an accurate value of lag phase and μmax, the maximum specific growth rate.

### 2.9. Enzymatic activity

To eliminate metabolites that could interfere with enzymatic activity measurements, the crude extract was desalted in Sephadex G-25 Columns (GE Healthcare) and reduced with 20 mM DTT in 50 mM Tris-HCl (pH 7.5) for 30 min at 30°C. The PRK activity was measured as previously described (Gurrieri et al., 2019). Briefly, the reaction mixture contained 50 mM Tris-HCl (pH 7.5), 1 mM EDTA, 40 mM KCl, 10 mM MgCl_2,_ 5 U/mL Pyruvate kinase, 6 U/mL Lactate dehydrogenase, 2.5 mM phosphoenolpyruvate, 2 mM ATP, and 0.2 mM NADH. The desalted crude extract was added and the background was recorded for 1-2 min. PRK activity was then measured by adding 0.5 mM ribulose-5-phosphate and monitoring the oxidation of NADH at 340 nm.

## 3. Results and discussion

### 3.1. The absence of PRK impacts the growth of Chlamydomonas

Phosphoribulokinase is an enzyme unique to the CBBC encoded by a single gene (Cre12.g554800_4532) in *Chlamydomonas reinhardtii*. To comprehensively investigate the function of the PRK-encoding gene, we used the mutant LMJ.RY0402.119555 (hereafter named ΔPRK) from the Clip Library, generated by random insertion of the paromomycin resistance gene (Li et al., 2019). This mutant strain harbors an insertion (named CIB1) in the exon 7 of the PRK gene (Figure 1A) as identified by junction sequencing with a 95% confidence (Li et al., 2019). In the mutant strain, the position of the CIB insertion in exon 7 was verified by PCR analysis with primer couples allowing specific detection of a single band related to either the WT PRK gene or the disrupted PRK gene containing the CIB1 insertion (Figure 1B). To determine the impact of the CIB1 insertion on PRK protein expression, total protein extracts from the mutant strain and the reference strain were analyzed by western blotting. The PRK signal was very strong in the reference strain, while no signal could be detected in the mutant strain (Figure 1C). To evaluate the detection threshold of our custom made anti-PRK polyclonal antibody, a gradient of total protein from the reference strain CC-4533 was utilized and a PRK signal could be still be detected using 5% of the total protein extract (Figure 1C).

**Figure 1:**
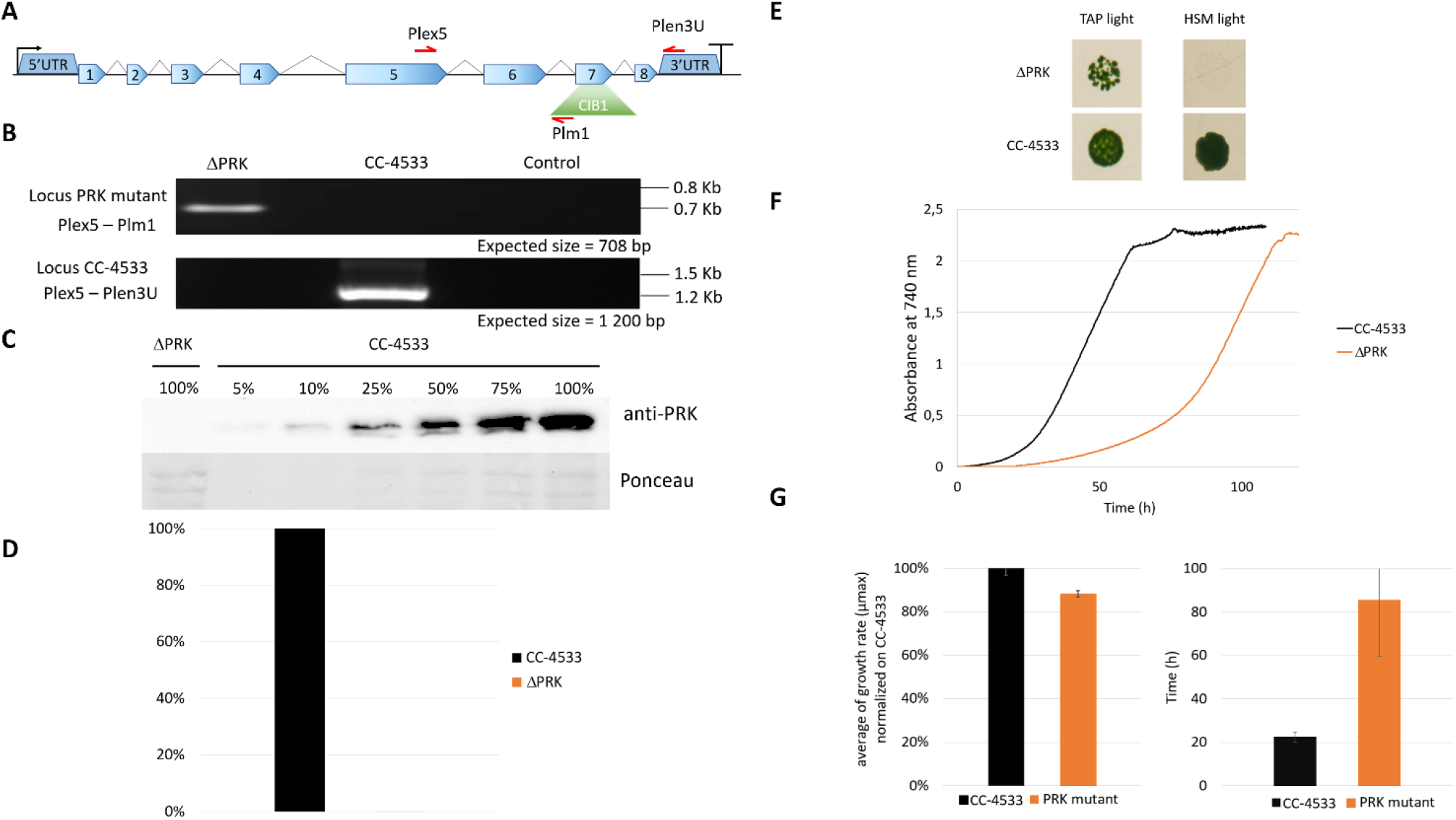
Characterization of the PRK mutant. (A) Representation of the PRK genomic locus in the Clip mutant with the CIB1 insertion (containing the paromomycin resistance gene) positioned in exon 7. (B) PCR on genomic DNA to confirm the presence of the CIB1 insertion in the PRK gene in the mutant strain (Locus PRK mutant) and the presence of the intact PRK locus (Locus CC-4533) in the reference strain. (C) Anti-PRK western blot on total soluble protein extracts of CC-4533 and ΔPRK strains (100% corresponds to 12 μg of total protein). (D) Total activity of reduced PRK. Desalted crude extracts of CC-4533 and ΔPRK were reduced with 20 mM DTT prior to activity measurement. (E) Spot test in TAP and minimal (HSM) media under continuous light (25°C, 100 μmol photons m^-2^ s^-1^). The reference strain CC-4533 was used as a control. 10^6^ cells were spotted and incubated for 7 days prior to observation. (F) Growth profile comparison between ΔPRK and CC-4533. Cultures were inoculated at 10^5^ cells/mL and incubated in TAP under light (25°C and 100 μmol photons m^-2^ s^-1^). (G) Kinetic parameters of growth for ΔPRK and CC-4533 strains, calculated from the data in panel F. Left graph: mean of the maximal growth rate (μ_max_), Right graph: mean of the lag phase duration. Error bars represent the standard deviation on a biological triplicate. B, C, E and F are one representative experiment out of 3 biological replicates.

This finding indicates that the mutant strain has a PRK content lower than 5% of that in CC-4533, strongly suggesting that the presence of the CIB insertion disrupts PRK protein expression. Since PRK activity is unique in Chlamydomonas, the absence of the PRK protein should correlate with a loss of the corresponding enzymatic activity. As PRK is activated by reduction (Marri et al., 2008), we treated the total soluble protein extract from each strain with the strong reducing agent DTT to obtain maximal PRK activity. Consistently, no PRK activity was detected in the mutant strain compared to the reference strain (Figure 1D). Consequently, the LMJ.RY0402.119555 strain is a knock-out strain with undetectable levels of both PRK protein and activity.

To examine the functional consequences of PRK deficiency in Chlamydomonas we analyzed the growth phenotype of the ΔPRK mutant under various conditions. Since PRK is central to the CBBC, we tested the ability of this mutant to grow in the presence of light and acetate as a carbon source (TAP light, mixotrophic conditions) or in the presence of light in a minimal medium without acetate (HSM light, strictly photoautotrophic conditions). In solid media, the ΔPRK mutant could not grow photoautotrophically but was able to grow in the presence of light and acetate, albeit more slowly than the reference strain (Figure 1E). To further characterize the growth phenotype of the ΔPRK strain in mixotrophic conditions, we monitored its growth over time in a photobioreactor in TAP liquid medium, compared with the CC-4533 reference strain (Figure 1F). The growth profiles revealed a pronounced difference between the two strains. Indeed, the mutant grew much slower than CC-4533, explaining the difference observed on solid media. This growth defect of the PRK mutant is characterized by an extended lag phase (Figure 1G) and a reduced maximum growth rate (μ_max_) compared to the reference strain (Figure 1G). These data indicate that the ΔPRK strain is non-photoautotrophic and strongly imply that the PRK gene is essential for photosynthetic carbon fixation, and that the absence of the CBBC limits the growth of the knock-out strain. To demonstrate that this phenotype is due to the absence of PRK, we sought to functionally complement the ΔPRK strain.

### 3.2. Complementation of the *PRK* mutant shows that PRK is essential for photoautotrophy

To verify whether the loss of photoautotrophic growth is due to the absence of PRK in the mutant, we functionally complemented the ΔPRK strain with a synthetic construct designed to restore PRK expression. This construct was built using the intron-less coding sequence (CDS) of the PRK gene, obtained from cDNA, fused to a triple HA tag under the control of the promoter, 5’UTR and 3’UTR/Terminator of PSAD gene. This transcriptional unit was coupled to a hygromycin resistance gene (Figure 2A). The PSAD genetic elements were previously shown to allow strong constitutive expression of a reporter gene (Crozet et al., 2018). The synthetic construct containing the two transcriptional units was introduced into Chlamydomonas nuclear genome by transformation. Multiple clones from the transformation were selected in the light on TAP solid medium supplemented with hygromycin and named C_x_, for complemented strain X. The C_x_ strains were genotyped by PCR using primer couples allowing specific detection of either the CIB insertion characteristic of the ΔPRK strain background or of the synthetic transgene (Figure 2B). All five chosen C_x_ and the mutant were found to contain the CIB1 insertion confirming the ΔPRK background. All selected clones also displayed the presence of the synthetic PRK transgene that was absent in the ΔPRK strain. We performed anti-PRK western blots to confirm the expression of the transgenic PRK. As shown in Figure 2C, 3xHA tagged PRK, which has a higher molecular weight than endogenous PRK, is present in all C_x_ complemented strains. In comparison with the gradient of CC-4533 total protein extract, we observed that the expression of the transgenic PRK-3HA is much lower than the endogenous PRK and shows some variability between C_x_ clones. This may be due to the design of the synthetic PRK construct and/or to position effects due to insertion of the transgene in distinct genomic sites.

**Figure 2:**
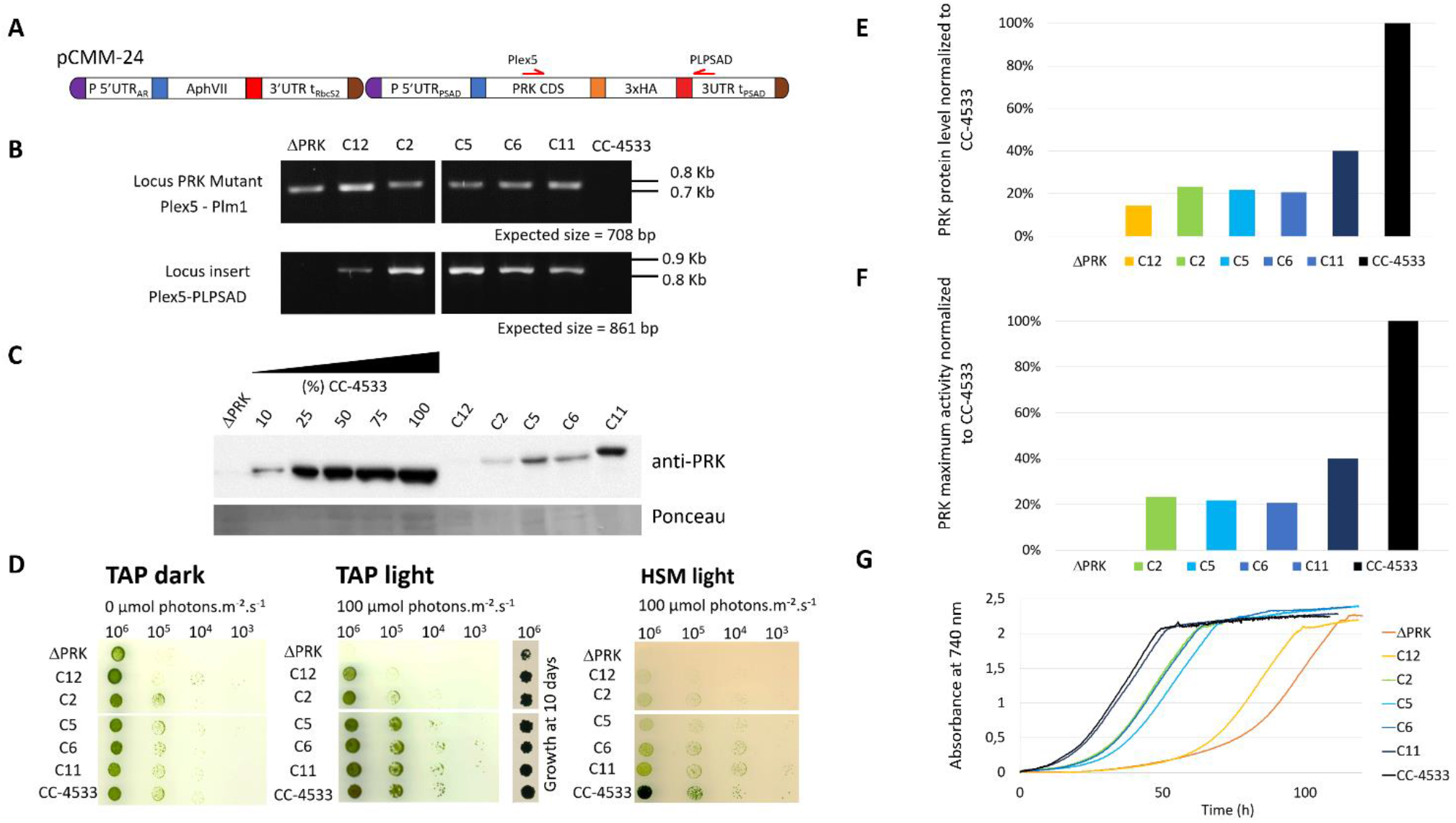
Functional complementation of the PRK mutant. (A) Design of the pCMM-24 construct used for PRK mutant complementation. (B) PCR on genomic DNA to confirm the presence of the CIB1 insertion in the PRK gene in the ΔPRK strain (Locus PRK mutant) and the presence of the pCMM-24 insertion in the complemented strains (Locus insert) in the complemented strains (C_x_ strains). (C) Anti-PRK western blot on total soluble protein extracts of the CC-4533, ΔPRK and C_x_ strains (100% corresponds to 12 μg of total protein). (D) Spot test in TAP and HSM minimal media after 5 days growth at 25°C in the dark or in the light (100 μmol photons m^-2^ s^-1^). The number of cells spotted is indicated above the spot test image. To show the growth of the ΔPRK strain in TAP light conditions, the growth after 10 days is shown. (E) Relative quantification of the PRK protein content in the C_x_ strains compared to CC-4533, calculated from the data in panel C. (F) Total activity of reduced PRK in the different strains. Desalted crude extracts of WT, C_x_ and ΔPRK were reduced with 20 mM DTT prior to activity measurement. (G) Growth profile of selected strains. Cultures were inoculated at 10^5^ cells/mL and incubated in TAP under light (25°C and 100 μmol photons m^-2^ s^-1^).

If the loss of PRK in the mutant is responsible for its growth phenotype, a functional complementation restoring photoautotrophic growth in minimal medium should be observed in the C_x_ strains. The growth phenotype of the different strains was assessed using a spot test analysis on solid media (Figure 2D).

In TAP medium in the dark, Chlamydomonas grows heterotrophically and all C_x_ strains grew like the reference strain. In TAP medium in the light, the complemented strains C_x_ grew similarly to CC-4533 although C_12_ appeared to grow significantly more slowly. By contrast the ΔPRK strain grew much more slowly than all other strains, as shown previously (Figure 1E). Nevertheless, after 10 days the growth the ΔPRK strain was clearly visible (Figure 2D). In HSM medium in the light, a restoration of the photoautotrophic growth was observed in the C_x_ strains. This demonstrates that PRK is essential for photoautotrophy in Chlamydomonas. However, the growth of C_x_ strains appeared slower compared to CC-4533 suggesting a partial functional complementation. To assess whether this phenotype was caused by a lower level of PRK protein, the relative quantity of PRK expressed in the complemented strains was determined. All the C_x_ strains showed a relatively low expression of the PRK enzyme, with a maximum of 40% of the PRK content compared to the reference strain (Figure 2E). The C_12_ strain that has a barely detectable level of PRK still partially restored photoautotrophy. PRK activity measurements fully correlated with the PRK levels estimated by western blot (Figure 2F). To quantitatively assess the growth limitation imposed by decreased PRK contents, we performed growth kinetics in a controlled photobioreactor. The C_x_ strains grew in TAP liquid medium in accordance with their PRK content (Figure 2G), as observed in the spot test assays (Figure 2D). Importantly, the strain with 40% of PRK content (C_11_) behaved as the reference strain in TAP medium but not in minimal medium (Figure 2D). This suggests that 40% of PRK content compared to the reference strain is limiting the CBBC in Chlamydomonas. In tobacco, the content of PRK leading to a phenotype was below 15% of the reference (Paul et al., 1995; Banks et al., 1999). Therefore, the excess of PRK may be more limited in Chlamydomonas than in tobacco. In order to determine more precisely the PRK content limiting the CBBC and to test whether overexpression of PRK may increase the CBBC turnover, a higher level of expression of the transgene needs to be achieved.

Although position effect may account for the variations observed between transformants, the limitation is more likely due to the design of the synthetic transgene (Figure 2A). Indeed, we screened hundreds of transformants and never found any clone with a PRK level above 40% of the level of the endogenous PRK, as exemplified in Figure 2. This suggested that the genetic elements controlling PRK expression may not be appropriate to ensure an expression level comparable to the reference strain.

### 3.3. New synthetic construct designs for enhanced PRK expression

The limitation to about 40% of the protein level found in the reference strain when expressing PRK from the synthetic construct may stem from various factors. PRK, like numerous other CBBC enzymes, is highly abundant and is estimated to represent 0.25% of total cellular proteins in Chlamydomonas (Hammel et al., 2020). Although the strong constitutive PSAD promoter and 5’UTR were employed to drive PRK expression in the synthetic pCMM-24 construct, it is plausible that the endogenous PRK promoter is significantly stronger and would ensure a higher expression level. The absence of introns in the synthetic gene may also be partly responsible for the low expression. Indeed, it is now well-established that the presence of introns in the synthetic construct is frequently required to ensure high-level expression of a transgene in Chlamydomonas (Lumbreras and Purton, 1998; Fuhrmann et al., 1999; Baier et al., 2018). The presence of introns could boost gene expression due to the presence of a transcriptional enhancer, through a process called Intron-mediated enhancement that stimulates directly transcription or through interaction with the spliceosome (Schroda, 2019; Baier et al., 2020). An online tool has been developed to design Chlamydomonas transgenes with artificial introns (Jaeger et al., 2019). The fusion of the recombinant PRK with a triple HA-tag may also impact expression, potentially by destabilizing the protein or affecting its activity.

To investigate the significance of the promoter strength and of the presence of introns we have designed new synthetic constructs that incorporate the endogenous PRK promoter and its 5’UTR to drive either expression of the PRK CDS devoid of introns and also lacking triple HA-tag or the native PRK coding sequence containing natural introns (Figure 3A). The PRK promoter and the native PRK gene coding sequences were amplified by PCR from Chlamydomonas CC-4533 genomic DNA and cloned as level 0 MoClo parts. These parts were then used to build the two transcriptional units later on coupled to the hygromycin resistance gene to generate pCMM-25 and pCMM-26 (Figure 3A). Each of these synthetic constructs were introduced in the nuclear genome of the ΔPRK strain. The original synthetic gene (pCMM-24) containing the PSAD promoter and PRK CDS was also transformed again in the ΔPRK strain as a control. The transformants were selected on photoautotrophic growth restoration in minimal medium. It was observed that more colonies were obtained for transformations with pCMM-26 than with pCMM-25 or pCMM-24, suggesting that the presence of introns had a positive effect on the number of functionally complemented strains. For each transformation, roughly 50 colonies were selected for further analysis.

**Figure 3.**
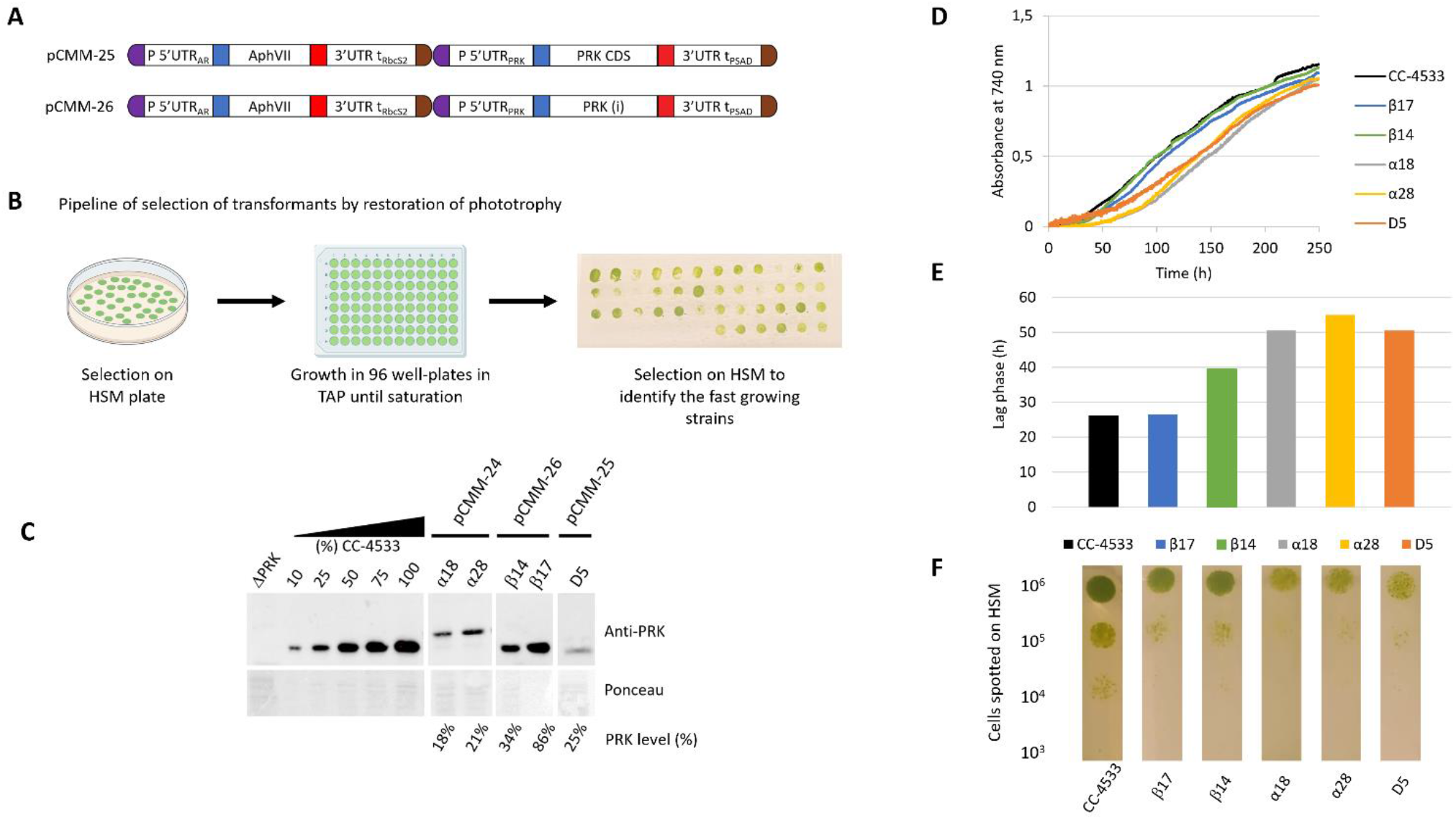
New designs to achieve higher PRK expression. (A) Design of the pCMM-25 and pCMM-26 constructs. (B) Pipeline for the identification of strains with high PRK content by selection of transformants on HSM plates, followed by growth to saturation in 96-well plates in TAP medium, and re-plating on HSM agar plates. The strains with the fastest growth on the plate were selected for further analysis. (C) Anti-PRK western blot on total soluble protein extracts of the ΔPRK, CC-4533, and complemented strains obtained by transforming the ΔPRK strain with the pCMM-24 (α strains), pCMM-26 (β strains) or pCMMM-25 (D5 strain) constructs. 100% corresponds to 12 μg of total protein. A gradient of the total protein extract of the strain CC-4533 was used for the quantification of the PRK content in the different complemented strains, indicated as PRK level (%). (D) Growth profile of CC-4533 and complemented strains. Cultures were inoculated at 10^5^ cells/mL and incubated in HSM at 25°C under continuous light (100 μmol photons m^-2^ s^-1^). (E) Lag phase measured for CC-4533 and complemented strains, calculated from the data in panel D. (F) Spot test on HSM minimal medium after 5 days of growth at 25°C in the light (50 μmol photons m^-2^ s^-1^). The number of cells spotted is indicated on the left and the strain is indicated below each spot test line.

As we previously observed with the first synthetic construct that in the ΔPRK background cell growth is directly correlated with PRK expression level (Figure 2), we selected the fastest growers to increase the probability to select strains with higher PRK levels through spot test assays on HSM medium after growth to saturation in TAP medium (Figure 3B). For several representative fast growers, we analyzed the PRK content by immunoblot (Figure 3C) and the growth properties in liquid HSM medium in a photobioreactor (Figure 3D, 3E) or in solid HSM medium (Figure 3F). As previously shown (Figure 2), the pCMM-24 construct (α strains) only allowed partial complementation of the ΔPRK phenotype. The level of PRK was limited to 21% in the best case and photoautotrophic growth was only partially restored compared to CC-4533. Comparable results were obtained with the pCMM-25 construct as exemplified by the best growing strain D5 which PRK level was limited to 25% of the reference strain CC-4533 and consistently showed partial complementation of photoautotrophic growth. This indicates that neither the expression without the triple-HA tag nor the use of the PRK promoter to drive PRK CDS expression resulted in increased PRK expression level. Conversely, the pCMM-26 construct expressing the intron-containing PRK endogenous coding sequence allowed to fully complement the phenotype of the ΔPRK strain (Figure 3) and even overexpress the protein (Figure 4). This indicates that the presence of introns in the PRK coding sequence is crucial for high level expression of the transgene. This result is consistent with previous studies (Lumbreras and Purton, 1998; Fuhrmann et al., 1999; Baier et al., 2018; Lauersen et al., 2018; Schroda, 2019; Baier et al., 2020). It would be interesting in future studies to determine whether the high-level expression is linked to a specific PRK intron. Even with the pCMM-26 intron containing construct, a range of PRK expression levels (34%-250% relative to CC-4533) was observed in the transformants. This variability is most likely attributable to random insertion of the construct in regions of the genome with a variable ability to drive gene expression. This position effect is classically observed in Chlamydomonas and can be exploited to generate strains with diverse levels of expression of a given transgene. In our case, immunoblot and growth assays revealed that a PRK content of 86% is sufficient to fully restore photoautotrophic growth with kinetics comparable to the reference strain CC-4533 (Figure 3). This result suggests that PRK may be present in excess in Chlamydomonas. Nevertheless, this excess appears much more limited than previously reported in tobacco where PRK limitation was only observed below 15% of the WT level (Paul et al., 1995; Banks et al., 1999). To confirm that PRK endogenous level does not limit the CBBC we analyzed strains overexpressing PRK.

**Figure 4.**
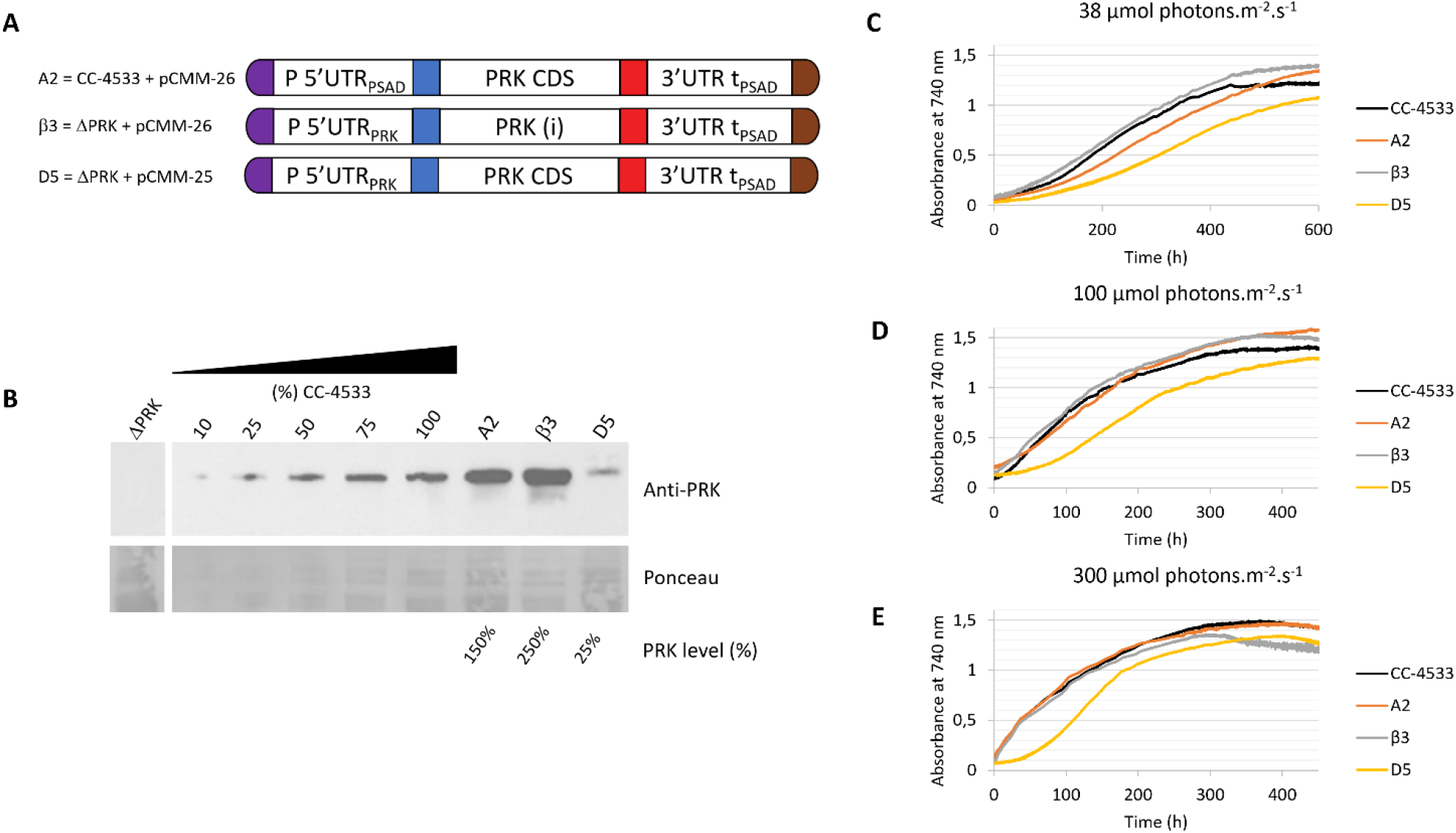
Overexpression of PRK. (A) Design of the transgene driving PRK expression and the genetic background (ΔPRK or CC-4533) for the A2, β3 and D5 strains (B) Anti-PRK western blot on total soluble protein extracts of the ΔPRK, CC-4533, and transformed strains as indicated. 100% corresponds to 12 μg of total protein. A gradient of the total protein extract of the strain CC-4533 was used for the quantification of the PRK content in the different complemented strains, indicated as PRK level (%). (C), (D), (E) Growth curves of CC-4533 and transformed strains. Cultures were inoculated at 10^5^ cells/mL and incubated in HSM at 25°C under three distinct light intensities as indicated.

### 3.4. Overexpression of PRK does not affect cell growth

To analyze the impact of PRK overexpression in Chlamydomonas, we examined strains transformed with the pCMM-26 construct either in the ΔPRK or the CC-4533 backgrounds (Figure 4A). The growth of strains overexpressing PRK 1.5-fold (A2 strain) or 2.5-fold (β3 strain) (Figure 4B) was analyzed in a photobioreactor in liquid HSM medium under three different light intensities (Figure 4C-E). Strains expressing 100% (CC-4533) or 25% (D5) of PRK were used as controls. The growth of the two overexpressor strains was comparable to the CC-4533 strain. At high light intensity (300 μmol m^-2^ s^-1^) growth of the overexpressor was similar to CC-4533 while at lower light intensities the 1.5-fold overexpressor showed a marginally slower growth kinetics compared to the two other strains. The cause of this slower growth in low light of the A2 strain is not known but could be related to the site of insertion of the transgene. Nonetheless, these results clearly show that overexpression of PRK does not lead to an increased growth. This suggests that the endogenous level of PRK in Chlamydomonas is not limiting the Calvin-Benson-Bassham cycle under optimal growth conditions.

## 4. Conclusions

In the present study, we have characterized a PRK knock-out mutant of Chlamydomonas and utilized this strain to investigate the limitations imposed by PRK expression level on the CBBC. We demonstrated that PRK is essential for photosynthesis in *C. reinhardtii* through a comprehensive analysis of the mutant and its functional complementation. The essentiality of PRK for photoautotrophic growth was previously suggested by a large-scale systematic characterization of gene function in Chlamydomonas (Fauser et al., 2022). Previously, the only analyses of PRK deficiency in Chlamydomonas were obtained in the F-60 mutant generated by chemical mutagenesis and without a full validation by functional complementation (Moll and Levine, 1970). This mutant was also shown to spontaneously revert ((Smith, 2000) and personal observation of the authors). Through our functional complementation approach, we demonstrated that the level of PRK protein needs to be as high as 86% of the level of the reference strain to allegedly restore the standard growth phenotype. Moreover, we established that, in the conditions tested, overexpression of PRK does not improve Chlamydomonas growth. This suggests that the endogenous PRK content is not limiting the CBBC in *Chlamydomonas reinhardtii*. PRK is therefore in excess in Chlamydomonas but this excess is much more limited than observed in tobacco where PRK limitation was only observed below 15% of the WT level (Paul et al., 1995; Banks et al., 1999). This implies that a slight decrease of PRK level or PRK activity would be sufficient to limit the CBBC in Chlamydomonas. This may allow PRK to play a more prominent role in the control of the CBBC turnover in Chlamydomonas, for example under conditions leading to light-dependent regulation of PRK activity mediated by thioredoxins and CP12 (Gurrieri et al., 2021; Gurrieri et al., 2023). Nevertheless, the CBBC functioning is different in algae compared to land plants, particularly because a carbon-concentration mechanism (CCM) in the pyrenoid increases carbon fixation by Rubisco (Barrett et al., 2021) and likely imposes a stronger requirement for RuBP production by PRK to sustain growth, especially in non-limiting light conditions. Metabolite profiling revealed that RuBP is indeed significantly more concentrated in algae and cyanobacteria compared to land plants (Clapero et al., 2023). This higher concentration of RuBP may be required to ensure that substantial concentration gradients drive rapid diffusion into the CCM compartment that houses Rubisco (Treves et al., 2022). Finally, the PRK knock-out strain along with the genetic elements we have generated may constitute useful tools to explore PRK regulation *in vivo* using functional complementation with PRK variants harboring targeted mutations such as mutations to serine/alanine of regulatory cysteines.

## 6. Author Contributions

NDB, GF, MM, MZ, SDL and PC designed the study and analyzed the data. NDB, GF, MM, MZ, CCC, SDL and PC discussed and wrote the manuscript. NDB, GF, MM, LJ and PC performed the experiments.

## 8. Acknowledgments

We thank Ferdinand Meneau, Dr Théo Le Moigne, Dr. Christophe Marchand, Dr Antoine Danon, and Dr. Julien Henri for stimulating discussions and suggestions.

